# Exact confidence intervals for population growth rate, longevity and generation time

**DOI:** 10.1101/2023.04.12.536246

**Authors:** Carlos M Hernandez-Suarez, Jorge Rabinovich

**Affiliations:** Universidad Francisco Gavidia; CEPAVE, CCT La Plata, CONICET- UNLP

**Keywords:** Confidence intervals, Generation time, Life history traits, Life table, Longevity, Population growth rate

## Abstract

By quantifying key life history parameters in populations, such as growth rate, longevity, and generation time, researchers and administrators can obtain valuable insights into its dynamics. Although point estimates of demographic parameters have been available since the inception of demography as a scientific discipline, the construction of confidence intervals has typically relied on approximations through series expansions or computationally intensive techniques. This study introduces the first mathematical expression for calculating confidence intervals for the aforementioned life history traits when individuals are unidentifiable and data are presented as a life table. The key finding is the accurate estimation of the confidence interval for *r*, the instantaneous growth rate, which is tested using Monte Carlo simulations with four arbitrary discrete distributions. In comparison to the bootstrap method, the proposed interval construction method proves more efficient, particularly for experiments with a total offspring size below 400. We discuss handling cases where data are organized in extended life tables or as a matrix of vital rates.

## Introduction

Estimating life history traits (LHTs), such as growth rate, longevity, and generation time, can provide critical insights into population dynamics. The finite growth rate, *λ*, is a particularly important parameter, as it measures the number of individuals produced per unit of time per individual. The *instantaneous growth rate, r*, is a related measure, where *λ* = *e*^*r*^. However, estimating the growth rate is usually made by following a group of individuals which introduces natural sampling errors into the life history trait estimates, highlighting the importance of calculating confidence intervals. Despite this being a substantial problem, there are currently no specific mathematical expressions for calculating confidence intervals for the growth rate that show their dependance with the sample size, as it should be.

There are two main approaches to constructing confidence intervals for the growth rate, namely the *series expansion* and the *numerical methods*, with the latter mostly based on resampling. Lande (1988) was the first to use the theory of series expansions, known in statistics as the *delta method*, to derive approximations to the variance of the growth rate *λ* (see Alvarez-Buylla and Slatkin 1991, 1994; Sibly et al. 2000). Series approaches assume that errors in estimating the vital rates are small, allowing for straightforward derivation of confidence intervals.

Matrix projection models, which include the Leslie matrix for age-classified populations and the Lefkovitch matrix for populations classified into arbitrary life stages, relate the number of individuals in a particular age or stage category at time *t*, **n**(*t*), to that at time *t* + 1 as **n**(*t* + 1) = **A n**(*t*), where **A** is a matrix containing vital rates per stage. In these models, the variance for the growth rate *λ*, can be approximated through a series expansion involving the estimated vital rates *a*_*ij*_, the elements of **A**. This method can be extended to estimate other LHTs.

Computer-intensive methods such as resampling, specifically bootstrap and jackknife, have been used to estimate confidence intervals for LHTs (Efron and Tibshirani 1986). Bootstrap involves resampling from the original sample and analyzing the variation in estimates, while jack-knife involves excluding observations. Both methods were once computationally demanding, but advances in computing power now allow for a large number of iterations to be performed quickly on most desktop computers. Bootstrap assumes that sampling with replacement from the original sample of size *n* mimics taking a sample of size *n* from a larger population. It has been widely used (Meyer et al. 1986) but it may not work well for small samples. Overall, both methods, series expansion and resampling approaches are effective for estimating confidence intervals for LHTs, with each method offering unique advantages and limitations depending on the size and characteristics of the sample.

Monte Carlo simulations are a class of computer-intensive methods used to estimate LHTs in population projection models. These methods assume that the distribution of the estimates is known, allowing for sampling from that distribution to obtain a new dataset that is then analyzed to obtain LHTs. For instance, in a matrix model for a stage-classified population, the fraction of individuals moving from stage *i* to stage *j* at the next unit of time can be represented as *a*_*ij*_. If there are **n**_*i*_(*t*) individuals in stage *i* at time *t*, a Binomial (**n**_*i*_(*t*), *a*_*ij*_) distribution can be used to simulate the number of individuals moving from *i* to *j* at time *t* + 1. All events that may occur at the next unit of time, such as transitions between stages, reproduction and deaths must be simulated, and the simulation stops at a specified time or when the population reaches an equilibrium, that may include extinction. Each simulation produces a series of LHTs, which are then used to infer statistical properties such as the mean, variance, and confidence intervals for these parameters (see Alvarez-Buylla and Slatkin 1993). The Monte Carlo simulation method can be computationally intensive, but it can be useful for situations where other methods are not applicable or practical.

In matrix projection models the growth rate *λ*, is estimated by finding the largest positive eigenvalue of **A**. For further details see Caswell (2001). In matrix models characterized by stage-classified populations, the matrix **A** can be decomposed as **A** = **U** + **F**, where **F** represents a matrix of fertilities and **U** is a matrix of stage transitions, the equivalent to the transpose of a Markov chain with transient states. Caswell (2009) derived the variance of certain life history traits using limiting distribution results for passage times, as delineated by Iosifescu (1980). For example, if **N** = (**I** − **U**)^−1^, the expected longevity *E*[*L*] is provided by the sum of the first column of the elements in the matrix **N**, and the variance of longevity is given by:

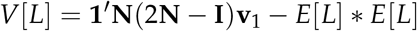

with **v**_1_ being a vector of zeros with a 1 in the first entry. Nevertheless, this and other expressions assume that the parameters of the transition matrix **U** have been estimated without error, a notion that is evident given the absence of *n*, the sample size, in the mathematical expressions. In this scenario, if **U** is treated as a Markov chain, the results on expectation and variances presented in Caswell (2009) do not differ from those that would be obtained using a Monte Carlo approach. Recall that the variance of a random variable *Y* that depends on an estimate of a parameter 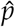 is

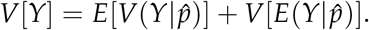

By assuming 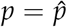, we are taking 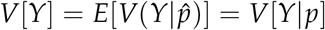 and neglecting the second term. In general, the demographic literature lacks comprehensive discussions on the statistical properties of estimates for the aforementioned life history traits. In the present study, we demonstrate how to derive point and interval estimates of longevity (*L*), generation time (*μ*_1_), and the instantaneous growth rate (*r*) using the statistical properties of these estimates.

In many cases, individuals within a population can be identified, enabling the measuring of individual age at reproduction, total offspring size per individual, and time until death. However, in certain stage-classified population analyses and models, individuals are not identifiable, allowing researchers only to count the number of individuals in each stage at time *t*. Newborns can be counted per time unit, but their mothers cannot be identified, a situation referred to as *anonymous reproduction*. The most limiting scenario arises when only the total number of living individuals and total newborns at each time unit are known. In this study, we address this specific situation where only three data columns are available: time units, number of living individuals, and offspring produced during each time unit. It is important to note that this represents the least informative scenario, as all other data organization (counts per stage or knowledge of offspring per individual at every unit of time) can be condensed to the data format utilized here.

In other words, the point and interval estimates proposed in this study can be applied to more complex forms of data organization beyond life tables.

Given these constraints, we are unable to provide variance estimates for *R*_0_, the *basic reproductive number*, which is considered one of the most critical parameters in demography. As we will demonstrate, *R*_0_ is the quotient of two easily observed quantities that measure the population’s reproductive capacity. However, our assumption of anonymous reproduction prevents the assignment of offspring to individuals and, consequently, the acquisition of measures of variation. Considering the close relationship between *r* and *R*_0_ highlighted in Wallinga and Lipsitch (2007), providing intervals for *r* serves as a means of addressing uncertainty regarding the population’s growth rate.

## Methodology

We will focus on discrete models, in which the different events (births, deaths) occur within a specified interval of unitary size. Throughout this study we employ vector notation, denoting a column vector as **x**, where *x*(*i*) represents the *i*-th element of **x**. The product **x**′**y** refers to the cross-product of two vectors, yielding a scalar. The element-by-element multiplication of two vectors is denoted by **x** ∘ **y**, which is also known as the Hadamard product and results in a vector. Time intervals are labeled 1, 2, 3, …. For each time *t*, we record the number of individuals alive, *n*(*t*) and the total offspring produced, *h*(*t*). In vector notation, we have three columns **t, n** and **h**. Using this initial information, we generate two additional columns: **g** and **f**. First, we define *K* = ∑ *h*(*i*) as the total offspring produced during the study period, and *N* as the initial number of individuals, which corresponds to *n*(1). The vector **g** is an estimate of the probability mass function of the time to reproduction, obtained by normalizing **h**, that is, **g** = **h**/*K*. The vector **f** contains elements *f* (*i*) = [*n*(*i*) − *n*(*i* + 1)]/*N*, representing the number of individuals that died in the *i*-th interval. Note that **f** is one element shorter than **n**, so we append a 0 at the bottom of **f** to maintain compatibility with the other arrays. Notice **f** is an estimate of the distribution of deaths.

To construct confidence intervals of size 1 − *α*, we will use a value *Z*_1−*α*/2_ representing the 1 − *α*/2 quantile of a standard Normal distribution. We will start from the traditional definition for each LHT.

### Longevity

The life history trait *Longevity* (*L*) is defined as the expected number of time units that an individual survives. It is usually calculated using the fraction of individuals that survive up to time *t* denoted by *l*(*t*), where *t* represents the time units:

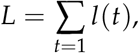

but *l*(*t*) = *n*(*t*)/*N*, where *n*(*t*) is the number of individuals alive at time *t* and *N* is the initial number of individuals. Thus, an estimate of *L*, denoted by 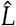, is given by:

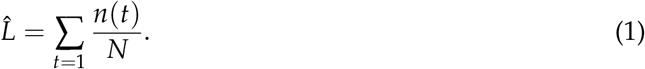

To calculate the variance of the estimate 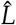, we can recalculate equation (1) in an alternative way. Let *X* be a random variable representing the time to death. Then, we have:

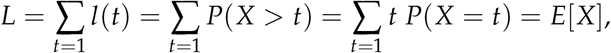

since *P*(*X* = *t*) is estimated with **f**, we can estimate the mean and variance of the longevity *L* using:

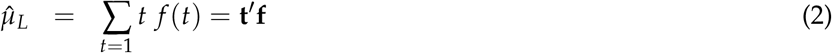

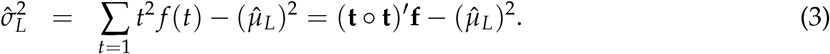

As we can see from (1), 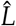 is an average and consequently, it follows asymptotically a Normal distribution with mean 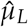 and variance 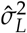. A confidence interval of size 1 − *α* for *L* is:

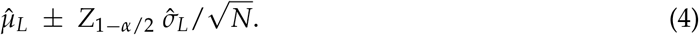

### R_0_, the Basic Reproductive Number

The *Basic reproductive number* (*R*_0_) also known as the *net reproductive net* is a fundamental quantity in population biology that measures the average number of offspring produced by an individual over its lifetime. The usual expression to calculate *R*_0_ is:

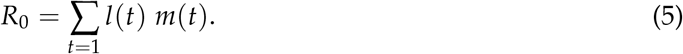

where *m*(*t*) is the fertility at time *t. R*_0_ is required to calculate the *Mean generation time*. However, we cannot provide the variance of our estimate of *R*_0_ unless some distributional assumptions (that we do not want to concede here) are made.

We can simplify equation (5) using again the fact that *l*(*t*) = *n*(*t*)/*N* and *m*(*t*) = *h*(*t*)/*n*(*t*). Thus, we have:

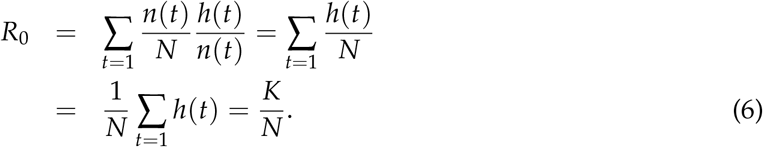

Since *K* is the total offspring produced in the study period and *N* is the initial number of individuals, *R*_0_ is the total offspring divided by the initial number of individuals. Strictly, since *N* is the initial number of individuals and these are usually newborns, *R*_0_ is the total number of newborns produced by a newborn. If an individual goes through a series of *M* stages namely *S*_1_, *S*_2_, …, *S*_*M*_, *R*_0_ could be defined as the average number of individuals of stage *j* produced by an individual of the same stage. Whatever *j* we choose, the result should be the same, although is easier to start with newborns.

Sometimes it is argued that since only females produce offspring, the analysis of *R*_0_ should be done considering females only. Equation (6) shows that this discussion is irrelevant. Nevertheless, it is preferable to include all individuals, males and females, since larger sample sizes are associated with smaller standard deviations. Furthermore, for some species, incorporating sex differentiation may introduce an additional layer of complexity that could potentially burden the analysis.

### The Generation Time μ_1_

The generation time, denoted by *μ*_1_, is defined as the average age of females at the time they bear offspring, considering that a mother can have multiple offspring at different ages. The generation time is usually calculated with the following expression:

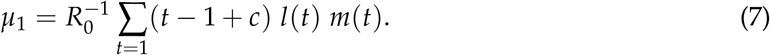

were *c* is the time within the interval where individuals are born, or instance *c* = 0 if births occur at the beginning and *c* = 1 if they occur at the end (Hernandez-Suarez 2011). In what follows we set *c* = 1.

By substituting *R*_0_ = *K*/*N*, we can rewrite (7) as:

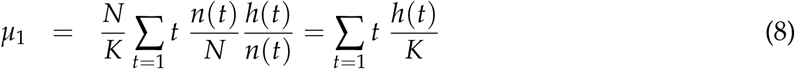

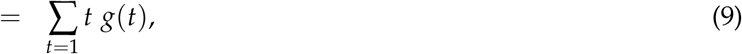

where *g*(*t*) represents the distribution of newborns at time *t*. To calculate the mean and variance of the generation time, we can define a random variable *T* as the average age at bearing as:

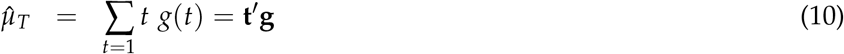

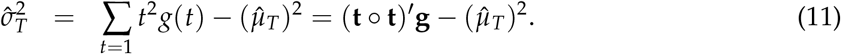

It is important to note that in equation (8), ∑ *t h*(*t*) represents the sum of the mothers’ ages at which each of the *K* offspring was produced. As a result, 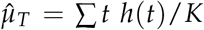 is an average over *K* observations, and subsequently, it follows asymptotically a Normal distribution with a mean of 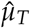 and a variance of 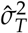. Therefore, a confidence interval with a size of 1 − *α* for *T* can be expressed as follows:

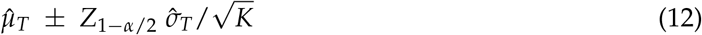

### The instantaneous rate of population growth, r

The Lotka-Euler equation relates growth rate with survival and fertility. It was initially formulated for continuous time by Lotka (1907a,b), while its discrete-time counterpart was developed by Cole (1954). We will use the discrete version, which corresponds to a life table. The *instantaneous growth rate r* is the value such that:

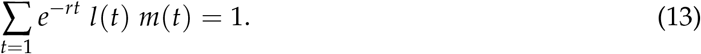

The growth rate *r* is found by numerically solving the previous expression. Recall *l*(*t*) is estimated with *n*(*t*)/*N* and *m*(*t*) with *h*(*t*)/*n*(*t*). Since *R*_0_ = *K*/*N*, substituting these values in (13) yields:

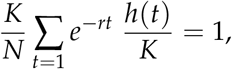

but *g*(*t*) = *h*(*t*)/*K* is the distribution of female age at offspring, and *K*/*N* = *R*_0_, thus, we arrive to the discrete time version of the Lotka-Euler equation:

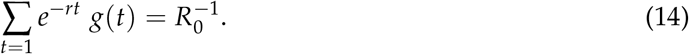

For the equivalent derivation for continuous time see Wallinga and Lipsitch (2007).

### Confidence intervals for r and λ

First we rewrite (14) as:

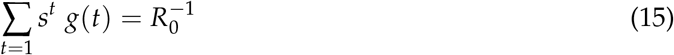

with *s* = *e*^−*r*^. The left side of (15) is the *probability generating function* (pgf) of *g*(*t*), that is, *E*[*s*^*t*^], provided that |*s*| ≤ 1. Clearly, for *r >* 0 this requirement is fulfilled. The situation *r <* 0 will be discussed later in this section.

The following result regarding probability generating functions is well known (see the review in Nakagawa and Perez-Abreu 1993):

Let *x*_1_, *x*_2_, *x*_3_, …, *x*_*n*_, be a random sample from a discrete distribution *F* over 0, 1, 2, …, with corresponding probabilities *p*_*k*_, *k* = 0, 1, 2, The *empirical probability generating function* (epgf) is defined as:

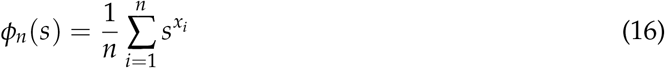

for |*s*| *<* 1. This transform of the empirical distribution function *F*_*n*_, is an estimator of

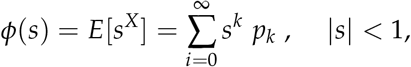

the pgf associated to *F*. For each fixed *s, ϕ*_*n*_(*s*) is an unbiased estimator of *ϕ*(*s*). Moreover, by the law of large numbers *ϕ*_*n*_(*s*) is a consistent estimator for *ϕ*(*s*), and by the central limit theorem

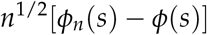

converges in distribution to a Gaussian random variable with zero mean and variance *σ*^2^(*s*) = *ϕ*(*s*^2^) − (*ϕ*_*n*_(*s*)) ^2^

We will use the previous results on epgf to derive confidence intervals for *r* in (14). Suppose that *p*_*k*_ is known, and that the solution of *s* in 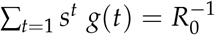 is *θ*. Let:

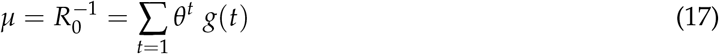

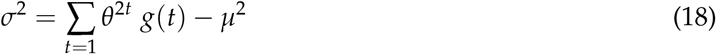

then

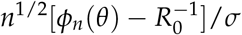

follows asymptotically a *N*(0, 1) distribution. In other words, for our fixed *θ*-value as the solution of (15), the epgf *ϕ*_*n*_(*θ*) tends to a Normal distribution with mean 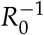 and variance *σ*^2^/*n*, with *σ*^2^ as in (18).

Now, let 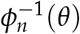 be the inverse function of *ϕ*_*n*_(*θ*). Observe that:

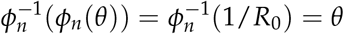

it follows that 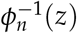 is the value of *s* such that

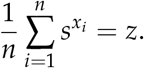

By the asymptotic properties of *ϕ*_*n*_(*θ*) we know that there are two values *L* and *U* such that:

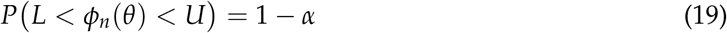

where

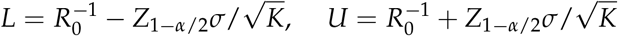

then from (19):

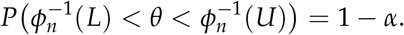

Then a CI of size 1 − *α* for *θ* is (*θ*_*L*_, *θ*_*U*_) where *θ*_*L*_ and *θ*_*U*_ are the solutions of:

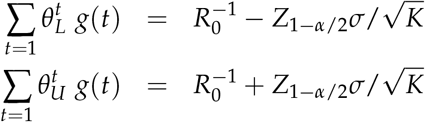

### Implementation of the epgf method

In practice, we use exp(−*r*) instead of *s* in (15), and the observed vector **g** as an estimate of *g*(*t*). We first estimate *R*_0_ using (6), then we estimate *r* with 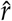, the solution for *s* in:

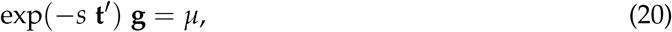

with *μ* = 1/*R*_0_. Then use this *r*-value in (18) to estimate *σ*^2^ with:

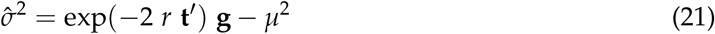

and finally derive confidence intervals as the solutions of *r*_*L*_ and *r*_*U*_ in:

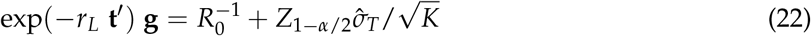

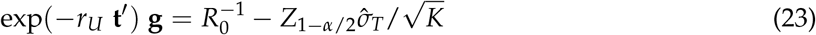

Similarly, point an interval estimates of the same size for *λ* are:

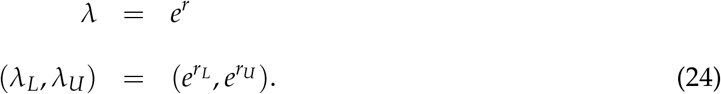

We develop a full exercise in the Appendix applying these formulae, but first, we will test the accuracy of the interval proposed with Monte Carlo simulations.

### Negative growth rates

It is important to note that the theory presented earlier, concerning epgf’s, is strictly valid only when *r >* 0, implying that it seems to be functional exclusively when the instantaneous growth rate *r* is greater than 0. But the condition *r >* 0 has been established to ensure that the sum in equation (15) converges to a finite value, as we must remember that function *g*(*t*) is defined for *t* = 1, 2, 3, …, however, in real-life experiments, we use a function *g*(*t*) defined for a finite set of *t*-values, thus the sum may converge for a broader range of *r*-values, including negative values. The advice is simple: if a numerical solution for *r* exists in equation (15) (which could be negative), then the sum has converged, and that solution serves as our estimate of *r*.

## Simulations

The primary objective of these simulations is to assess the accuracy of expressions (22) and (23) by applying them to four arbitrarily selected discrete distributions that are described below, representing the offspring distribution. For each distribution, we first calculate point and interval estimates of *r* using Sec. where **g** is the known distribution. Then we conducted 10^5^ simulations, with each simulation *j* executed as follows:

1. Sample *n* values from the target distribution and obtain *x*_1_, *x*_2_, *x*_3_, …, *x*_*n*_. Call this sample *j*.
2. Compute 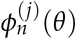 using (16) for this sample *j*.
3. Determine the solution for *s* in (20) for this sample and denote it as *s*(*j*).
4. Estimate the growth *r* for this sample as *r*(*j*) = − log(*s*(*j*)).

When the simulations were concluded, we calculated the mean and variance of 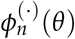, which are equivalent to *μ* and *σ*^2^/*n*, respectively. We also counted the fraction of the *r*(*j*)’s falling inside the theoretical CI for *r* given in (22) and (23).

### Distributions used

The following four distributions were used in the simulations:

1. Distribution 1. A Shifted Poisson distribution where *P*(*Y* = *y*) = *P*(*X* = *y* − 20), with *X* being a Poisson random variable with parameter of 7. In this case, *Y* = 20, 21, 22, …, (Fig. 1a).

**Figure 1:**
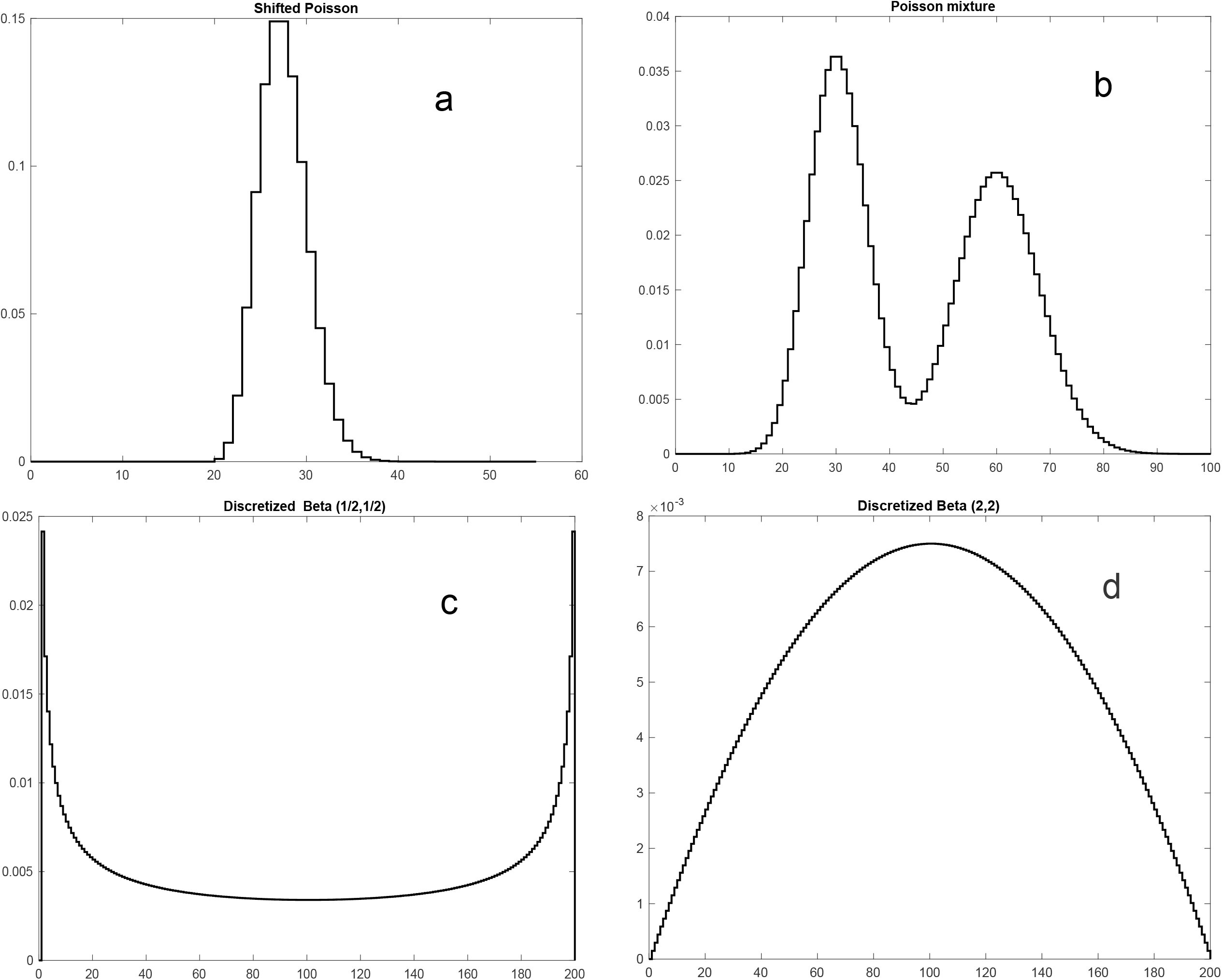
Offspring distributions *g*(*t*) used in the Monte Carlo simulations described in Section “Simulations”.
2. Distribution 2. Poisson mixture. Two Poisson random variables *X* and *Y* were generated with respective parameters of 30 and 60. The resulting distribution is bimodal, and the random variable *W* is defined such that *P*(*W* = *k*) = (*P*(*X* = *k*) + *P*(*Y* = *k*))/2, (Fig. 1b).
3. Distribution 3. Discretized Beta. For *t* = 0, 1, 2, …, 200, we calculated the probability mass function (pmf) of the random variable *Y* as *P*(*Y* = *t*) = *P*(*X* = *t*/200)/*c*, where *X* is a Beta (1/2, 1/2) distribution and 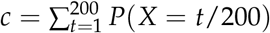, (Fig. 1c).
4. Distribution 4. Similar to the previous distribution, but with *X* being a Beta (2, 2) distribution, (Fig. 1d).

In all distributions, the value of *R*_0_ in equation (15) is arbitrarily selected as a constant for testing purposes.

## Results

In this study, Monte Carlo simulations were not employed to evaluate the confidence intervals for longevity and mean generation time provided in (4) and (12) because of the robust and unambiguous statistical foundations underlying the distribution of the sample mean. Instead, due to the less familiar nature of the empirical probability generating function *ϕ*_*n*_(*θ*), we focused our efforts on testing the confidence intervals for the growth rate.

The distributions employed for the age of mothers at reproduction in our simulations may be questionable, particularly distributions 2 and 3, but our objective was to test the estimates in situations that include distributions far from bell-shaped. From Table (1) and Figs (2a-2d), it can be observed that the distribution of the epgf *ϕ*_*n*_(*t*) closely aligns with a Normal distribution, with parameters as predicted. It is worth noting that we did not use large sample sizes in all cases, which could have increased the similarity between observed and expected distributions. Table (2) demonstrates that the fraction of the 10^5^ simulations falling inside the confidence interval closely matches the expected outcome across all tested distributions, for *α* = 0.01, 0.05 and 0.1

**Table 1:**
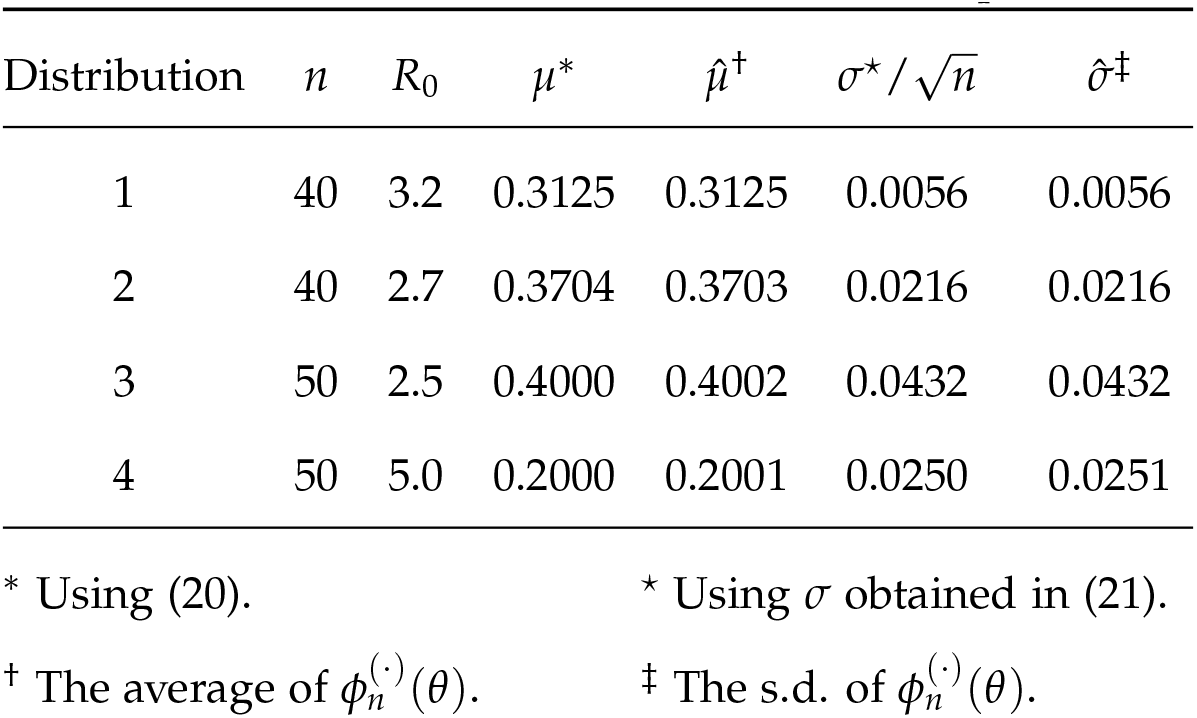
Comparison of first two moments of the theoretical and empirical distributions for *ϕ*_*n*_(*θ*)

**Figure 2:**
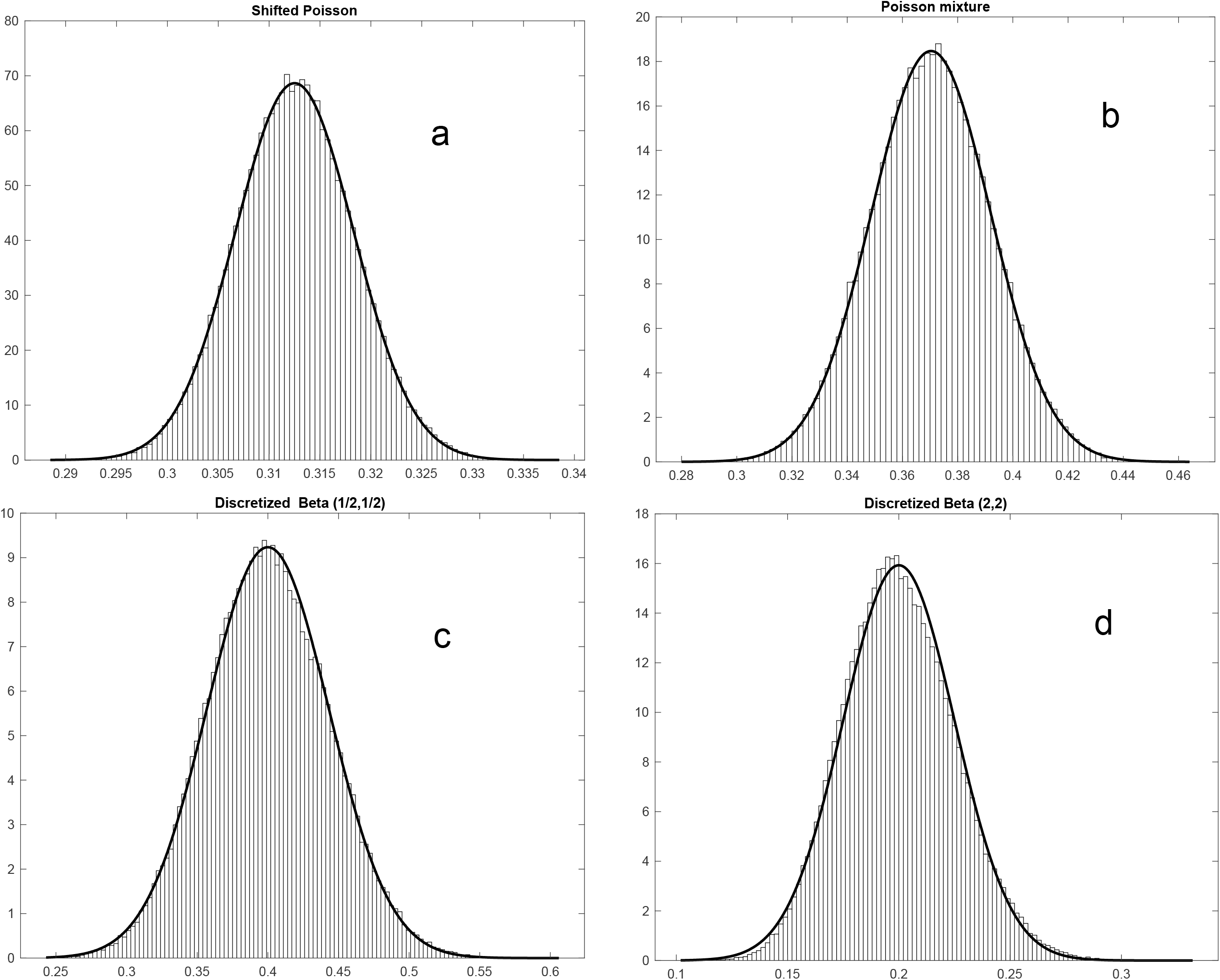
The histograms illustrate the observed distribution of *ϕ*_*n*_(*θ*) for each offspring distribution described in Section “Simulations’, using *θ* as the solution for *s* in (16). The superimposed distribution represents the theoretical (Normal) distribution, with mean and variance specified in (17) and (18). A comparison of the first two moments for the theoretical and empirical distributions of *ϕ*_*n*_(*θ*) is presented in Table 1.

**Table 2:** Theoretical (*r*^*^) and empirical 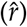 values of the growth rate. The table shows the CI_1−*α*_ and the fraction of samples where the estimated *r* was inside the predicted interval for *α* = 0.1, 0.05 and 0.01. Results of 10^5^ simulations as described in Sec..

| $1 - \alpha$ | Dist. | $n$ | $r^*$ | $\hat{r}^\dagger$ | $r_L^*$ | $r_U^*$ | $p^\ddagger$ |
| --- | --- | --- | --- | --- | --- | --- | --- |
| 0.90 | 1 | 40 | 0.043 | 0.043 | 0.042 | 0.044 | 0.899 |
|  | 2 | 60 | 0.023 | 0.023 | 0.021 | 0.026 | 0.900 |
|  | 3 | 40 | 0.012 | 0.012 | 0.009 | 0.016 | 0.902 |
|  | 4 | 50 | 0.019 | 0.019 | 0.017 | 0.023 | 0.902 |
| 0.95 | 1 | 40 | 0.043 | 0.043 | 0.042 | 0.044 | 0.949 |
|  | 2 | 60 | 0.023 | 0.023 | 0.020 | 0.026 | 0.951 |
|  | 3 | 40 | 0.012 | 0.012 | 0.009 | 0.018 | 0.952 |
|  | 4 | 50 | 0.019 | 0.019 | 0.016 | 0.024 | 0.952 |
| 0.99 | 1 | 40 | 0.043 | 0.043 | 0.041 | 0.045 | 0.989 |
|  | 2 | 60 | 0.023 | 0.023 | 0.020 | 0.027 | 0.990 |
|  | 3 | 40 | 0.012 | 0.012 | 0.008 | 0.020 | 0.989 |
|  | 4 | 50 | 0.019 | 0.020 | 0.015 | 0.026 | 0.991 |

\* Using (20).
$^\dagger$ The average of $r(j)$ 's as described in step 4 in Section "Simulations".
\* Obtained using (22) and (23).
$^\ddagger$ The fraction of the $r(j)$ 's that falls within the $CI_{1-\alpha}$ .

We also analyzed how confidence intervals (CI) obtained with bootstrap methods compare with the method introduced here. For this, we perform another set of simulations: for each distribution tested we took *n*_1_ = 10^4^ samples of size *n*, then each sample was resampled *n*_2_ = 10^3^ times to calculate the empirical CI_0.95_, and then recorded the fraction of the *n*_1_ intervals that contained the known value of *r*, which was recorded as *p*(*i, n*) since it is a function of the distribution used (*i* = 1, 2, 3, 4) and the value of *n*, for *n* = (10, 20, 30, 40, 50, 100, 200, 300, 400).

Figure 3 shows *p*_*n*_, the fraction of the simulations in which the predicted interval contained the true growth rate *r* as a function of sample size, averaging across all four distributions using bootstrap and the method proposed here based on epgf’s. The performance of bootstrap methods improves when the sample size *n* increases and is efficient for moderate *n*; however, it is less effective for small sample sizes. In contrast, the empirical probability mass function (empf), despite being founded on asymptotic properties of the sample, demonstrates greater efficiency for small sample sizes. The performance of the empf starts slightly above the threshold and exhibits faster improvement than the bootstrap method, approaching the anticipated 95% threshold by *n* = 30.

**Figure 3:**
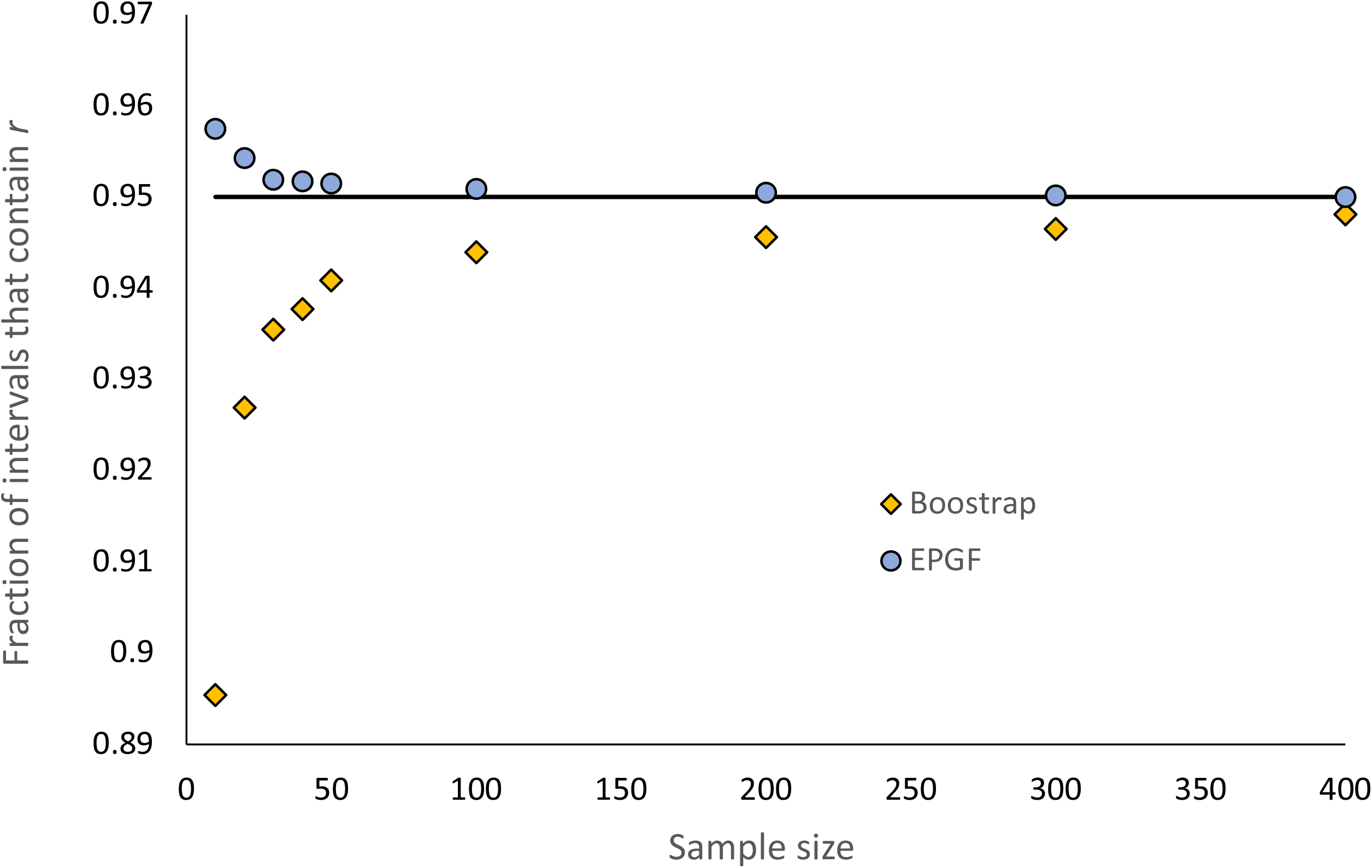
Comparsion with Boostrap. The plot shows the fraction of the intervals that contained the true growth rate *r* at different sample sizes for the Boostrap method (diamonds) and the epgf method (circles) proposed here. The horizontal line is the 0.95 target value. Convergence of Bootrsap is much slower. Details on the simulations are given in Section “Results”.

It is important to point out that in this context, the sample size *n* corresponds to the total number of available observations, which is the total offspring size *K*. Recall that distribution **g**, represents the offspring distribution, constructed using *K* observations.

## Discussion

It is interesting how the distribution of empirical probability generating functions (epgf) was absent from the construction of confidence intervals for *r*, despite the clear relationship between pgf’s and *R*_0_ given in equation (14). The confidence intervals provided in (22-23) are easily calculated from data organized as a life table and possess a solid theoretical framework.

Constructing confidence intervals for *R*_0_ cannot be achieved without making distributional assumptions. Nonetheless, given the close relationship between *R*_0_ and *r* shown in equation (14), the uncertainty regarding population growth is addressed through the confidence intervals for *r* or *λ*.

When the data is organized in a Leftkovitch matrix containing vital rates, including transition and fertility rates, which can be decomposed as **U** + **F**, we must recall that matrix **U** is generally obtained by estimating the likelihood of individual transitions from stage *i* to stage *j* at the next unit of time (for details see Hernandez-Suarez et al. 2019). If the original data that allowed the construction of **U** is available, the results derived here can be applied after collapsing the number of individuals alive at time *t* in all stages into a single column.

It is feasible to apply the proposed calculations to historical data contained in a life table with the traditional columns *t, l*(*t*), and *m*(*t*), as previously described if *N* or *K* is known. To begin, we can calculate *R*_0_ using equation (5), then, we obtain the columns **g** and **f** using the following equations:

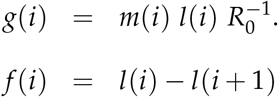

Furthermore, constructing confidence intervals requires the knowledge of both *N* and *K*, the total offspring size, to be incorporated into equations (4), (12), (22) and (23). Since *R*_0_ = *K*/*N*, knowing any of *N* and *K* will enable the calculation of the other.

In most entomological or fish demographic studies, with usually large offspring sizes, the bootstrap method will perform very well, although in some mammals or birds, with limited offspring, it would have limited use. Still, expressions (22-23) are more accurate since they involve the sample size. The method proposed here, based on properties of epgf’s, seems to perform very well for *K* ≥ 20, but in the simulations *p*_(*i,n*)_ never was below the 0.95 threshold even for *K* = 10.

## Conclusions

The proposed methodology for constructing confidence intervals has demonstrated efficacy, obviating the need for approximations through series expansion or reliance on computationally demanding techniques. While the method is specifically designed for data structured as life tables, it can be effortlessly adapted to extended life tables where the number of individuals at each stage is documented, or to data presented as a Leftkovitch matrix. It is worth mentioning that when the total offspring size (*K*) gathered in the experiment is below 400, the epgf approach demonstrates superior performance in comparison to the bootstrap method. https://github.com/car-git/Confidence-Intervals-LHT

## Appendix A: A complete example

This is an example of calculating point and confidence intervals for longevity (*L*), generation time (*μ*_1_), and growth rate (*r* and *λ*) using data from the study in Mermer et al. (2023).

The Brown marmorated stink bug, *Halyomorpha halys* (Staåal) (Hemiptera: Pentatomidae), is an invasive pest of over 300 plant species in North America, Asia, and Europe. It was first recorded in the US in 1996 and is a destructive agricultural and nuisance pest. It has high reproductive and dispersal capabilities and feeds on plant tissues, causing necrosis and scarring. The impact of temperature on its development, fecundity, survival, and mortality has been studied to determine its invasive potential and seasonal risk. Age-specific fecundity and survival data related to temperature extremes are needed to estimate reproduction, survival, and population dynamics.

This is part of the data set corresponding to the study in Oregon at 25°*C*. The data is shown in Table A1.

The only data required are columns *t, n*(*t*) and *h*(*t*), the rest is calculated as:

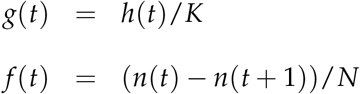

with *K* = 2430, the total egg production and *N* = 50, the initial number of individuals. A zero must be appended to the end of **f** to match the required length.

### Longevity

Using (2) and (3):

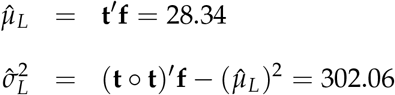

And the 95% confidence interval for *L* given in (4) are:

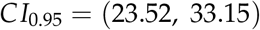

### *R*_0_

Using (6) we have:

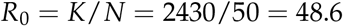

### Generation Time μ_1_

Using (10) and (11):

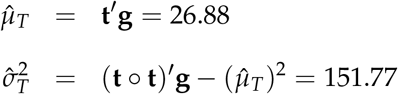

And the 95% confidence interval for *T* given in (12) are:

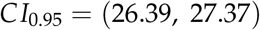

### The instantaneous rate of population growth, r

We need to solve for *r* in equation (20):

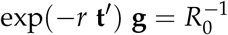

which yields *r* = 0.2021. To derive the lower limit of a 95% CI for *r*, we first use (21) to calculate 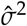, which is 7.1149 × 10^−4^, from here, 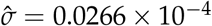

To derive the lower limit for *r*, we use equation (22) and solve numerically for *r*_*L*_ in:

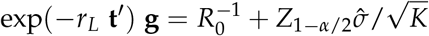

and the upper limit for *r* is found using equation (23) with the solution for *r*_*L*_ in:

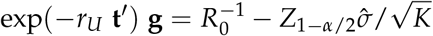

Thus the 95% confidence interval for *r* is (0.1988, 0.2055).

For *λ* we use 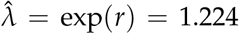 and the interval is (1.220, 1.228) is found as described in equation (24).

## Table for appendix

**Table A1:** Surving individuals **n** and number of eggs laid **h** by day **t**. Columns **g** and **f**, containing the relative frequency of eggs laid per day and mortality at time **t** are obtained as described in Section “Methodology”.

| **t** | **n** | **h** | **g** | **f** | **t** | **n** | **h** | **g** | **f** | **t** | **n** | **h** | **g** | **f** |
| --- | --- | --- | --- | --- | --- | --- | --- | --- | --- | --- | --- | --- | --- | --- |
| 1 | 50 | 0 | 0.0000 | 0.0000 | 26 | 23 | 72 | 0.0296 | 0.0400 | 51 | 6 | 0 | 0.0000 | 0.0000 |
| 2 | 50 | 0 | 0.0000 | 0.0000 | 27 | 21 | 42 | 0.0173 | 0.0200 | 52 | 6 | 0 | 0.0000 | 0.0200 |
| 3 | 50 | 0 | 0.0000 | 0.0200 | 28 | 20 | 57 | 0.0235 | 0.0000 | 53 | 5 | 0 | 0.0000 | 0.0000 |
| 4 | 49 | 0 | 0.0000 | 0.0000 | 29 | 20 | 94 | 0.0387 | 0.0400 | 54 | 5 | 0 | 0.0000 | 0.0000 |
| 5 | 49 | 0 | 0.0000 | 0.0000 | 30 | 18 | 71 | 0.0292 | 0.0200 | 55 | 5 | 0 | 0.0000 | 0.0200 |
| 6 | 49 | 0 | 0.0000 | 0.0000 | 31 | 17 | 77 | 0.0317 | 0.0000 | 56 | 4 | 0 | 0.0000 | 0.0000 |
| 7 | 49 | 0 | 0.0000 | 0.0200 | 32 | 17 | 25 | 0.0103 | 0.0000 | 57 | 4 | 3 | 0.0012 | 0.0000 |
| 8 | 48 | 0 | 0.0000 | 0.0400 | 33 | 17 | 30 | 0.0123 | 0.0200 | 58 | 4 | 22 | 0.0091 | 0.0000 |
| 9 | 46 | 0 | 0.0000 | 0.0400 | 34 | 16 | 0 | 0.0000 | 0.0400 | 59 | 4 | 28 | 0.0115 | 0.0000 |
| 10 | 44 | 28 | 0.0115 | 0.0000 | 35 | 14 | 76 | 0.0313 | 0.0000 | 60 | 4 | 0 | 0.0000 | 0.0200 |
| 11 | 44 | 28 | 0.0115 | 0.0000 | 36 | 14 | 68 | 0.0280 | 0.0000 | 61 | 3 | 0 | 0.0000 | 0.0000 |
| 12 | 44 | 10 | 0.0041 | 0.0200 | 37 | 14 | 43 | 0.0177 | 0.0000 | 62 | 3 | 0 | 0.0000 | 0.0000 |
| 13 | 43 | 118 | 0.0486 | 0.0400 | 38 | 14 | 77 | 0.0317 | 0.0200 | 63 | 3 | 14 | 0.0058 | 0.0000 |
| 14 | 41 | 111 | 0.0457 | 0.0000 | 39 | 13 | 82 | 0.0337 | 0.0200 | 64 | 3 | 0 | 0.0000 | 0.0000 |
| 15 | 41 | 136 | 0.0560 | 0.0400 | 40 | 12 | 4 | 0.0016 | 0.0400 | 65 | 3 | 0 | 0.0000 | 0.0000 |
| 16 | 39 | 162 | 0.0667 | 0.0400 | 41 | 10 | 61 | 0.0251 | 0.0200 | 66 | 3 | 0 | 0.0000 | 0.0000 |
| 17 | 37 | 110 | 0.0453 | 0.0400 | 42 | 9 | 0 | 0.0000 | 0.0000 | 67 | 3 | 0 | 0.0000 | 0.0000 |
| 18 | 35 | 49 | 0.0202 | 0.0400 | 43 | 9 | 0 | 0.0000 | 0.0000 | 68 | 3 | 0 | 0.0000 | 0.0000 |
| 19 | 33 | 231 | 0.0951 | 0.0400 | 44 | 9 | 29 | 0.0119 | 0.0000 | 69 | 3 | 0 | 0.0000 | 0.0000 |
| 20 | 31 | 0 | 0.0000 | 0.0600 | 45 | 9 | 27 | 0.0111 | 0.0000 | 70 | 3 | 0 | 0.0000 | 0.0000 |
| 21 | 28 | 0 | 0.0000 | 0.0200 | 46 | 9 | 20 | 0.0082 | 0.0000 | 71 | 3 | 0 | 0.0000 | 0.0000 |
| 22 | 27 | 61 | 0.0251 | 0.0600 | 47 | 9 | 58 | 0.0239 | 0.0400 | 72 | 3 | 20 | 0.0082 | 0.0200 |
| 23 | 24 | 155 | 0.0638 | 0.0000 | 48 | 7 | 3 | 0.0012 | 0.0200 | 73 | 2 | 4 | 0.0016 | 0.0400 |
| 24 | 24 | 60 | 0.0247 | 0.0000 | 49 | 6 | 0 | 0.0000 | 0.0000 | 74 | 0 | 0 | 0.0000 | 0.0000 |
| 25 | 24 | 36 | 0.0148 | 0.0200 | 50 | 6 | 28 | 0.0115 | 0.0000 |  |  |  |  |  |

